# The effects of sequence length and composition of random sequence peptides on the growth of *E. coli* cells

**DOI:** 10.1101/2021.11.22.469569

**Authors:** R. Johana Fajardo C., Diethard Tautz

## Abstract

We study the potential for the *de novo* evolution of genes from random nucleotide sequences using libraries of *E. coli* expressing random sequence peptides. We assess the effects of such peptides on cell growth by monitoring frequency changes of individual clones in a complex library through four serial passages. Using a new analysis pipeline that allows to trace peptides of all lengths, we find that over half of the peptides have consistent effects on cell growth. Across nine different experiments, around 16 % of clones increase in frequency and 36 % decrease, with some variation between individual experiments. Shorter peptides (8–20 residues), are more likely to increase in frequency, longer ones are more likely to decrease. GC content, amino acid composition, intrinsic disorder and aggregation propensity show slightly different patterns between peptide groups. Sequences that increase in frequency tend to be more disordered with lower aggregation propensity. This coincides with the observation that young genes with more disordered structures are better tolerated in genomes. Our data indicate that random sequences can be a source of evolutionary innovation, since a large fraction of them are well tolerated by the cells or can provide a growth advantage.

## 1. Introduction

New genes can arise by two alternative mechanisms (Andersson, Jerlstrom-Hultqvist, & Nasvall, 2015; Chen, Krinsky, & Long, 2013; McLysaght & Guerzoni, 2015; Schloetterer, 2015; Tautz & Domazet-Loso, 2011; Van Oss & Carvunis, 2019). The first is through duplication and/or recombination of existing genes or gene fragments, which later accumulate mutations that render them different from their parental genes. The second is *de novo* evolution from previously non-coding sequences. While this was long thought to be unlikely, there is now plenty of evidence that the process has probably been active throughout evolution (James et al., 2021; Neme & Tautz, 2013; Neme & Tautz, 2016; Pavesi, Magiorkinis, & Karlin, 2013; Ruiz-Orera, Messeguer, Subirana, & Alba, 2014; Wilson, Foy, Neme, & Masel, 2017). However, since it is difficult to distinguish *de novo* evolution from duplication followed by divergence beyond sequence recognition (Weisman, Murray, & Eddy, 2020), one can prove true *de novo* evolution only for relatively recent events, where evolutionary time has not been enough for accumulation of too many mutations (Tautz & Domazet-Loso, 2011). Several dedicated studies on individual genes, including functional analyses, have been published (Cai, Zhao, Jiang, & Wang, 2008; Heinen, Staubach, Haeming, & Tautz, 2009; D. Li et al., 2010; Reinhardt et al., 2013; Xie et al., 2019). In addition to this, there are well-documented cases of peptides with biological function derived from randomly synthesized sequences (Bao, Clancy, Carvalho, Elliott, & Folta, 2017; Keefe & Szostak, 2001; Knopp et al., 2021; Knopp et al., 2019; Stepanov & Fox, 2007). Overall genome comparisons between recently separated species have suggested that *de novo* evolved genes arise continuously with a high rate, but can also get lost at high rates (Durand et al., 2019; Neme & Tautz, 2014; Palmieri, Kosiol, & Schlotterer, 2014; Zhao, Saelao, Jones, & Begun, 2014). This dynamic transformation of non-coding sequences into coding ones is very clear, especially in eukaryotes, where large parts of the non-coding genome are transcribed. Comparisons between closely related mouse populations and species revealed the transcription of these non-coding regions is subject to fast evolutionary change, such that within a time span of 10 million years the whole genome can become transcribed and thus subjected to evolutionary testing (Neme & Tautz, 2016). Hence, the raw material for *de novo* evolution, namely transcripts from initially non-coding DNA regions, is abundantly present.

Based on these insights, we previously developed an experimental approach to ask which fraction of random sequences has a potential biological function that could become subject to further adaptive evolution (Neme, Amador, Yildirim, McConnell, & Tautz, 2017). We expressed a library of sequences with random sequence composition in bacterial cells and monitored which sequences could provide a growth advantage or disadvantage to the cell in the context of four growth cycles of the whole library. The general experimental design for this experiment is shown in Figure 1.

**Figure 1.**
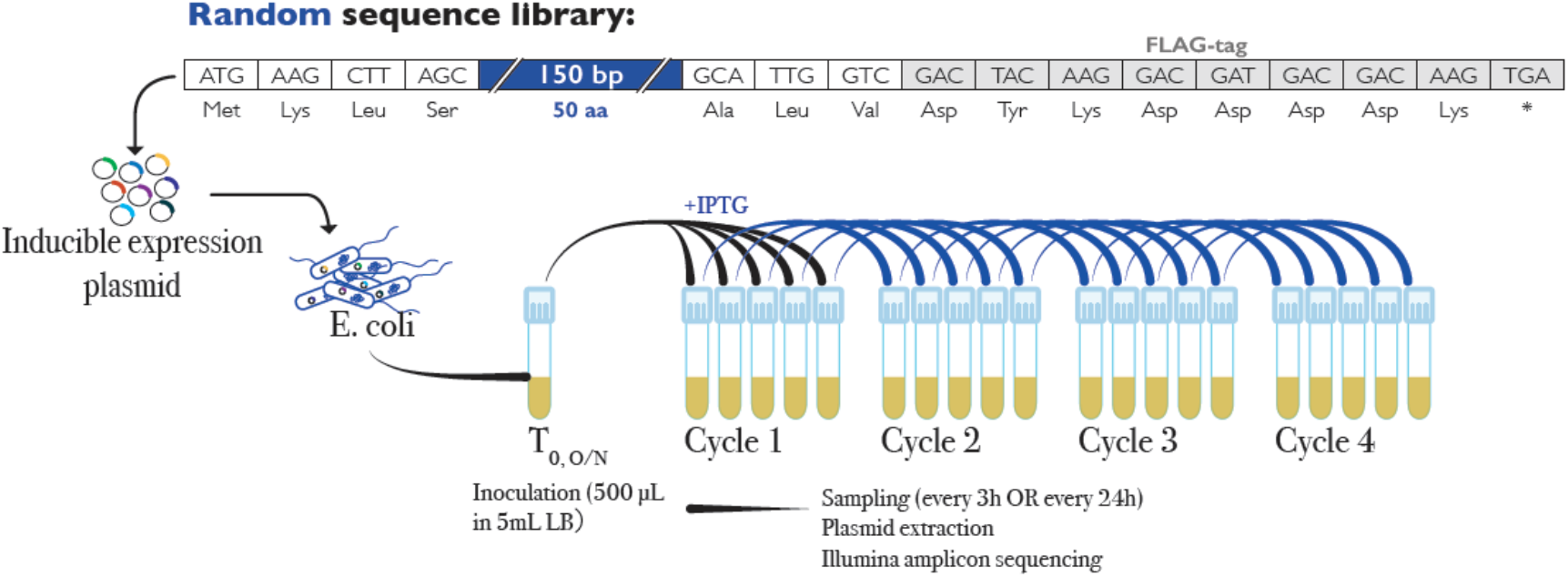
Experimental design to evaluate the fraction of bioactive sequences in a library of random sequences. A pool of random 150bp sequences generated by adding equimolar amounts of each nucleotide at every synthesis step was ligated into a commercial inducible expression vector (pFLAG-CTC, Sigma). This vector has start and stop codons in frame of the restriction site used for cloning, which means that the random sequences were flanked by a common sequence of 12 bp on the 5’-end and 36 bp on the 3’-end with a FLAG-tag (grey boxes). The resulting 195 nucleotide and 65 amino acid full sequences are shown. The pool of clones was used to transfect *E. coli* (DH10B) to generate a library of bacterial cells. Expression of the cloned peptides was induced by adding isopropyl β-d-1-thiogalactopyranoside (IPTG) to the culture media. Replicates were sampled every three hours for a total of 12 hours (3-hour cycles, 12-hour experiments) whereby one tenth of the culture volume was used for seeding the culture at each passage. The overall experiment replicated the one described in (Neme et al., 2017), where the analysis focused on full-length peptides only and included also experiments with 24-hour growth cycles (5-day experiments). Here we use a newly designed pipeline to analyse all experiments and all peptide lengths.

The experiments showed that a surprisingly large fraction of random sequences affected cell growth, either by enhancing it, or by slowing it down. In the initial analysis, between 11 to 25 % of the sequences increased in frequency in all replicates of each experiment, whereas 18 to 53 %, decreased (Neme et al., 2017). However, the study focussed exclusively on the full-length peptides in the library, although the design strategy with random synthesis of the insert produces also a large number of truncated peptides with premature stop codons.

In the present study, we first reproduced the experiment, but with a lower concentration of starting library in an attempt to reduce the possible impact of very many low-frequency clones on the overall mean fitness of the complex library. Plus, we designed a new pipeline to analyse the new experiment, as well as all of the previous experiments. This new pipeline allowed us to include the clones expressing truncated peptides and to assess whether the expressed vector without insert could have a growth effect on the cells harbouring it.

The new goal of this project was to explore the possible effects of shorter peptides in relation to the full-length peptides studied before. All peptides in the original analysis have common C-terminal residues (FLAG-tag—see Figure 1), which may have contributed to their stability and/or biological effects. Since naturally *de novo* evolved peptides would not have such a common C-terminus, it is important to verify whether the same spectrum of effects is also seen with peptides that have random C-termini. Furthermore, we wanted to explore sequence features of the peptides that could make them more or less likely to be tolerated by the cells and to be maintained in the population through several cycles of growth. Finally, we wanted to address the critical points that were raised against our original experiment, where (Weisman & Eddy, 2017) and (Knopp & Andersson, 2018) suggested a vector effect driving the patterns of peptides that rise in frequency. In this view, the vector itself would have a negative effect due to expressing a 38 amino acid peptide (or a secondary RNA structure) under induction conditions, which would be relieved when a “neutral” random sequence was replacing it, giving the impression that the “neutral” sequence acts positively. While we had argued that this effect could not fully explain the data that we had at that time (Tautz & Neme, 2018), further analysis of this question is certainly warranted.

## 2. Materials and Methods

### Library and replication experiment

We used the original library described in (Neme et al., 2017) from a stock frozen in 20 % glycerol. The general design of the library and the experiment are depicted in Figure 1. In order to assess whether there could be a complexity effect, we repeated the original experiment using a 100-fold dilution of the original library and a 1-day sampling schedule, with samplings every 3 hours for a total of 4 samplings in 12 hours. This was done by seeding 5 μL from the stock on 25 mL LB liquid medium with 500 μg/mL ampicillin, and allowed it to grow overnight at 37 °C with constant shaking (250 rpm). After 16 hours, 500 μL of the liquid culture were transferred into five 5 mL tubes containing 4.5 mL of LB medium with 10^−3^ mol/L IPTG to induce expression of the random sequences. For each cycle, 500 μL of culture from each tube were used to seed a new replicate after 3 hours of growth (37 °C, 250 rpm). From the remaining bacterial culture for each replicate, 3 mL were collected and used for plasmid extraction using a QI-Aprep Spin Miniprep kit (QIAGEN). Extracted plasmids were eluted in 30 μL of elution buffer and stored at −20 °C until use.

Amplicon sequencing of the library was performed using specific barcoded primers to amplify a 356-nucleotide fragment including the random sequences in a one-step PCR using PHUSION HF master mix (Invitrogen). The cycling program consisted of an initial denaturation at 98 °C for 30 seconds, followed by 25 cycles of 98 °C for 10 seconds, 65 °C for 20 seconds, and 72 °C for 1 minute. After a final elongation step of 72 °C for 10 minutes, samples were purified using a Qiagen MinElute Gel Extraction kit. Concentration of samples was calculated through relative quantification in an agarose gel, using a Molecular Imager(R) Gel Doc(TM) XR+ System with the Image Lab(TM) Software (Bio-Rad). Barcoded samples were pooled together in equal concentrations to obtain the sequencing library. Sequencing was done using Illumina’s MiSeq Reagent Kit v3 with 300 cycles to get overlapping 300-nucleotide paired-end reads.

### Available data

In addition to sequencing data from the diluted library experiment described above, we used the original fastq files for eight experiments described in (Neme et al., 2017). The original experiments were done following two different sampling schedules: either a 1-day course with samplings every 3 hours, or a 4-day course with samplings every 24 hours. In either case, four timepoints were sampled. The number of replicates, cycle duration and experiment length for each of the experiments are summarized in Table 1. In addition to three experiments with 10 replicates of each type of sampling schedule, we used two 4-day experiments with 5 replicates. One of them (experiment 7) was done with a treatment control without induction with IPTG, while the other one (experiment 8) was sequenced more deeply (5x more reads than the other experiments) to capture even rare clones present at low frequencies in the population.

**Table 1:** Clone performance in different experiments

| Exp<sup>1</sup> | Cycle length/ Experiment length (replicates) | N<sup>2</sup> | POS<sup>3</sup> | NEG<sup>3</sup> | NS<sup>3</sup> | Range log2-fold change (average) | Empty vector log2-fold change<sup>4</sup> |
| --- | --- | --- | --- | --- | --- | --- | --- |
| 1 | 3h / 1 day (n=10) | 5625 | 0.11 | 0.36 | 0.53 | -8.0 to 2.7 (-1.1) | 1.1 |
| 2 | 3h / 1 day (n=10) | 5606 | 0.17 | 0.43 | 0.41 | -7.7 to 2.2 (-1.2) | 0.5 |
| 3 | 3h / 1 day (n=10) | 5638 | 0.18 | 0.40 | 0.42 | -7.8 to 2.7 (-1.1) | 0.4 |
| 4 | 24h / 4 days (n=8) | 5623 | 0.14 | 0.30 | 0.56 | -5.2 to 5.2 (-0.5) | 0.1 |
| 5 | 24h / 4 days (n=10) | 5596 | 0.10 | 0.26 | 0.64 | -5.4 to 5.0 (-0.6) | -1.7 |
| 6 | 24h / 4 days (n=10) | 5632 | 0.26 | 0.41 | 0.32 | -5.9 to 2.2 (-0.7) | -1.1 |
| 7 | 24h / 4 days (n=5) | 5623 | 0.07 | 0.28 | 0.65 | -7.2 to 4.0 (-0.9) | -0.2 |
| 8 | 24h / 4 days (n=5) | 5689 | 0.27 | 0.46 | 0.27 | -11.2 to 1.4 (-1.6) | -0.4 |
| 9 | 3h / 1 day (n=5) / diluted library | 5651 | 0.16 | 0.32 | 0.51 | -8.4 to 5.6 (-0.7) | -0.7 |
| All experiments averages<sup>5</sup>: |  |  |  |  |  |  |  |
| All clones |  | 5621 | 0.16 | 0.36 | 0.48 |  |  |
| Clones with 4aa ORF |  | 200 | 0.04 | 0.73 | 0.18 |  |  |
| Clones with 5aa ORF |  | 221 | 0.02 | 0.52 | 0.44 |  |  |
| Clones with 6aa ORF |  | 209 | 0.06 | 0.37 | 0.56 |  |  |
| Clones with FLAG sequence |  | 638 | 0.03 | 0.77 | 0.17 |  |  |
| Clones with FLAG+1 sequence |  | 129 | 0.03 | 0.68 | 0.20 |  |  |
| Clones with FLAG+2 sequence |  | 126 | 0.05 | 0.64 | 0.23 |  |  |
| Clones 48+ aa without FLAG |  | 237 | 0.06 | 0.55 | 0.34 |  |  |

<sup>1</sup>For experiments 1- 8 we reanalysed the original fastq data from (Neme et al., 2017), experiment 9 with a diluted starting library was done within the framework of this study.
<sup>2</sup>Number of clones detected among the 5,701 unique sequence clones in the database for which at least 5 reads were mapped in each experiment.
<sup>3</sup>Fraction of clones in each category. POS and NEG were assigned when $p_{adj} < 0.05$ , otherwise the clone was categorized as non-significant (NS).
<sup>4</sup>All vector clone frequency changes were highly significant ( $p_{adj} < 0.01$ ) in their respective experiment, except for Exp 4 ( $p_{adj} > 0.05$ ).
<sup>5</sup>The distribution of values from the 9 experiments is not significantly different from a normal distribution (Shapiro-Wilk test, $p > 0.5$ ).

### Analysis Pipeline

First, the paired end reads for each experiment were trimmed using Trimmomatic (v. 0.36), and merged using the software USEARCH10 (-fastq_mergepairs, -fastq_maxdiffs 30, -fastq_minmergelen 100) (Edgar, 2010). Since each read in a pair covers the entire random sequence, up to 30 mismatches were allowed between the paired forward and reverse reads. The fastq_mergepairs algorithm resolves discrepancies between the forward and reverse reads by comparing the quality score for the conflicting position in each read. It keeps the residue with the best quality score in the merged read. Merging the reads with this algorithm reduces the percentage of sequencing errors kept in each read. Note that it is not possible to account for PCR errors that have occurred during the library preparation.

To remove reads that do not belong to a PCR product from the plasmids in the library, a custom Perl script was used to find and save all merged reads containing pre-defined sequences up- and downstream of the random sequences on the pFLAG-CTC plasmid. The pre-defined sequences were a 18bp sequence around the start codon, and the FLAG-tag, including the stop codon. The reads thus selected are considered clean amplicon reads, trimmed around the pre-defined sequences, and used for all subsequent analyses.

### Database generation

To generate a database of all unique sequences in the library that could be detected by the amplicon sequencing approach, all clean reads from all available experiments and replicates were first dereplicated using USEARCH10. Dereplication was done in 3 rounds. In the first round, the nucleotide sequences were sorted alphabetically, and the -fastx_uniques option was used to remove duplicate sequences, keeping only one sequence of each type in the database while keeping track of the number of total sequences of each type with the -sizeout option. In this way repeated identical sequences were removed and a “size” annotation was added to the read name indicating how many identical matches were present in the clean read files. In the second round, all files with singletons removed were merged into a single file of all amplicon sequences available, sorted and de-replicated again using the same exact-match method. This exact matching approach is prone to enrichment of PCR or sequencing errors, since any two reads with even a single nucleotide difference are kept as individual sequences in the database. Singleton reads–more likely to be PCR or sequencing errors–were removed and a third dereplication round using a clustering approach was implemented.

The third round of dereplication aimed to remove reads generated by PCR or sequencing artefacts. The clustering approach used is based on the one used for OTU validation in microbiome analyses. Reads were sorted in decreasing order of size annotation, and the -cluster_smallmem option of USEARCH10 was used with an identity cut-off of 0.97. The clustering algorithm used by USEARCH is a greedy clustering approach. Here, sorting by the size annotation means that high-frequency reads are used as centroids or seeds for clusters first. This strategy relies on the assumption that reads found in high frequencies are more likely to be real, and less-common, highly-similar reads are probably generated through PCR or sequencing errors. The identity threshold of 0.97 allows less frequent reads with, for example, up to 5 mismatches in the expected 195-nucleotide sequence to join the high-frequency centroids forming the clusters. Using an additional filter of minimum cluster size of 8 reads, commonly used in microbiome amplicon sequencing analyses, removes other artefacts from the database. The resulting library of unique clusters (full database, SuppData_BACT.tsv) was used as the final database.

This database served also basis for the simulation of a 100.000 sequence library in R by sampling A, T, G and C using the calculated probabilities for each nucleotide at each position (see below).

### Sequence features

Several parameters were used to characterise the sequences in the complete database, as well as in the sequence groups generated after mapping of the reads to find changes in frequency. ORFs were predicted using the program getorf form the EMBOSS suite (Rice, Longden, & Bleasby, 2000) using the full database as input (-minsize 12, -find 3). Only the first ORF was kept for each sequence. Predicted ORFs were translated in the first frame using transeq from the EMBOSS suite, and the first predicted peptide was kept for each ORF. Sequence length was calculated for each read, as well as the predicted ORF and peptide using bash programs.

The number of peptides of each length depends on the probability of getting a stop codon at each consecutive position, and not before. This is best described by the probability function of a geometric distribution:

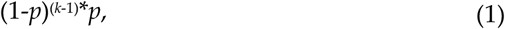

where *k* is the number of trials, in this case, the number of positions or the length of the sequence; and *p* is the probability of “success” or getting a stop codon. Multiplying this probability distribution by the number of synthesised sequences, one gets the expected count of peptides of each length. The resulting expected distribution of peptide lengths was used to confirm library quality (Supplementary Figure 2).

GC content was calculated as the percentage of guanine (G) and cytosine (C) in a sequence relative to its length using custom Perl scripts. This was done for the complete read, the random part of the sequence (obtained by trimming 12 nucleotides on the 5’-end and 33 nucleotides from the 3’-end of the clean reads), and the predicted ORF. Amino acid composition of the database and different sequence groups were calculated using the Biostrings package (V 2.58.0) from Bioconductor in R. Lists of sequences from each database formatted as AAStringSets were used as input for the letterfrequency function and amino acid frequencies were plotted for each sequence correcting for length. For the complete database, full-length predicted peptides were used, and frequencies were calculated for each sequence independently in order to obtain frequency distributions. For the group analysis, the flanking sequences were trimmed from the peptides and amino acid frequencies were calculated for the complete set of random amino acids as a single sequence.

Intrinsic disorder was calculated using the command line version of IUPred (IUPred2A) (Meszaros, Erdos, & Dosztanyi, 2018) with the -short option. Intrinsic disorder scores were averaged for each peptide to obtain single average disorder values. In addition to this, the fraction of residues with a predicted disorder score equal to or larger than 0.5 was calculated, producing comparable results (data not shown).

Protein aggregation propensity was calculated for all sequences in the database using the program PASTA 2.0 on the web server of The BioComputing POS lab of the University of Padua (Italy) (http://old.protein.bio.unipd.it/pasta2/) (I. Walsh, Seno, Tosatto, & Trovato, 2014). For each sequence, free energy for the single best pairing was obtained using the default settings for peptides. The best energy pairing for self-aggregation was obtained for each sequence from the output files, and energies of −5 or less were considered indicative of a high probability of aggregation.

### Mapping of reads to full database

Clean reads for all replicates and timepoints in each experiment were mapped to the database using a global alignment-based method from the program USEARCH10 (option -usearch_global). For consistency with the clustering analysis, alignments had a minimum required identity of 0.97, minimum query coverage of 0.9, and maximum one hit and 5 gaps. Hits were extracted from the search results and counted using custom bash scripts to generate count tables for each replicate in each experiment.

### Frequency change determination and group assignment

Raw count tables for each experiment were used as input for statistical analyses using the package DESeq2 in R (Love, Huber, & Anders, 2014). Count data of each experiment were analysed independently using cycle number as explanatory variable, and keeping only sequences that had at least 5 reads mapped in the whole experiment.

DESeq2 was designed mostly for the analysis of RNASeq data, but is broadly applicable to a large range of data types that require to control for large dynamic range and dispersion effects (Love et al., 2014). This makes different experiments better comparable between each other. Based on the log2-fold changes provided by DESeq2 (SuppData_DESeq2_ALLexp_Cycle4vs1.tsv) we classify the clones into NEG for negative changes and POS for positive changes. In addition, we chose the multiple-testing corrected padj value (provided by the program) as a cut-off to create a category of NS (“non-significant”) clones. For category assignment, a flag was added to each sequence on the database table depending on whether its fold-change was positive (> 1) or negative (< 1) and significant (p_adj_ < 0.05), or non-significant (p_adj_ > 0.05) for each experiment. For the overall assignments of sequences to one of the three categories, category flags were compared across all experiments, and a general flag (sign.most, in the database table) was assigned when at least the strict majority of experiments had the same flag (5 or more).

While P-values should normally not be used for a ranking between experiments, we believe that errors created in this way are small, or at least smaller than the variances that we see between the experiments anyway. A possible alternative for ranking the clones would be to calculate their individual fitness effects in the background of the mean fitness of the whole library, as suggested by (F. F. Li, Salit, & Levy, 2018). However, these authors advise against using their procedure under conditions where fitness distribution are broadly spread, as it is the case in our experiments (compare Figures 1 and 2 in (Neme et al., 2017)).

**Figure 2:**
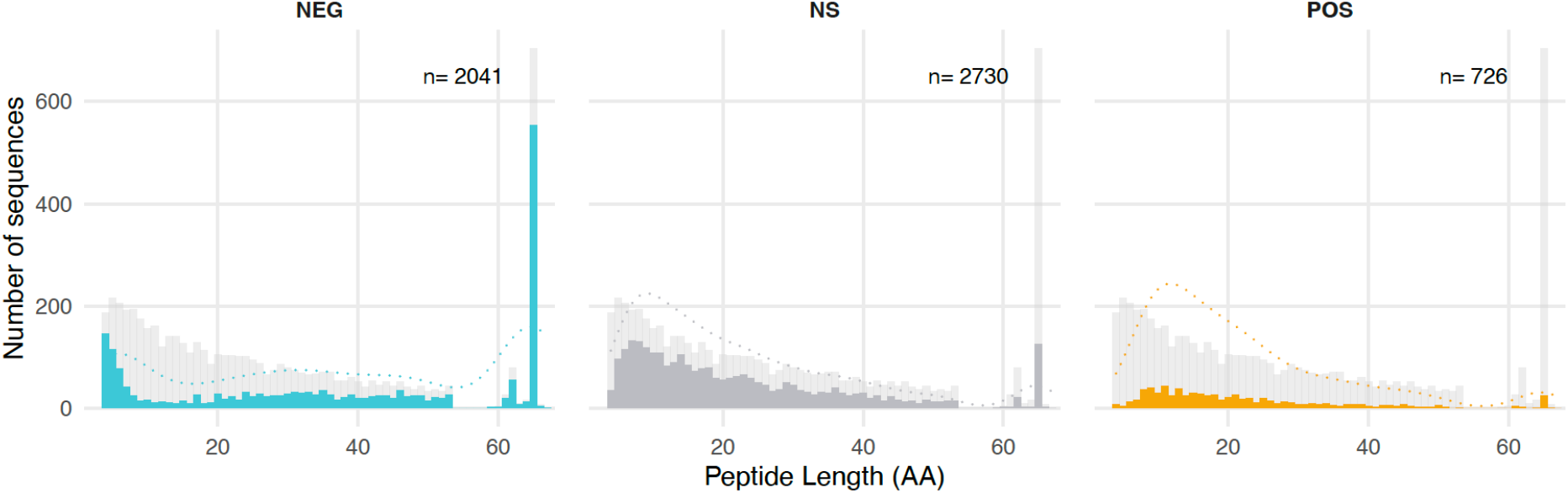
Length distribution of all predicted peptides in the random sequence database and assignment to response groups. Histogram of sequence lengths for each group of sequences. The coloured bars represent the number of peptides of each length assigned to each group in the experiments. Light blue: peptides showing a decrease in frequency (NEG); dark grey: peptides showing no significant change in frequency (NS); orange: peptides showing an increase in frequency (POS). The light grey bars in each panel represent the predicted peptide lengths of the complete database (compare suppl. Figure 2). Dashed lines represent the kernel density estimates for each category.

## 3. Results

### Replication with diluted library

In experiments with a complex library, all clones compete against each other, but rare clones generate only few reads that cannot be reliably analysed. Hence, these unaccounted background clones could influence the behaviour of the more frequent clones. In attempt to test this possibility, we repeated the experiment of (Neme et al., 2017), but with a starting stock that was diluted by 100-fold compared to the previous ones and used a sampling schedule with samplings every 3 hours for a total of 4 samplings in 12 hours. The further experimental steps were conducted as described in (Neme et al., 2017). The overall results showed that there was no major difference compared to the previous results (see Table 1 below). The majority of clones identified in the previous experiments could again be detected even with a 100-fold dilution. Hence, we decided to do the in-depth analysis described below across all available data.

### Characterization of the sequences in the random clone library

To analyse all experiments done with the given clone library, we first produced a reference sequence database including all different sequences reliably detected in any of the sequencing experiments. This required the establishment of a pipeline for filtering of PCR and sequencing errors, which we conducted based on a common approach that is also used in microbiome studies. We required that each sequence was represented by at least eight reads, biasing against rare variants that can be generated in the PCR amplification steps before sequencing.

The median number of paired-end reads per replicate was 284,875. On average, 79.3 % of them could be successfully merged, and both known plasmid-derived sequence regions could be found in 96.44 % (±1.46 %) of those merged. The resulting database consisted of 5,701 unique sequences with minimum cluster size of 8. This includes 647 peptides with the FLAG-tag sequence, of which 25 are not full-length due to internal deletions. 253 peptides end with frameshift versions of the FLAG-tag sequence. Furthermore, since for the random part of peptides of lengths 4, 5 and 6 there are only 1, 21, and 441 possible different amino acid sequences, respectively, different clones can code for the same peptide. For example, the library includes 200 clones coding for the shortest possible peptide (MKLS - derived from the vector, see Figure 1), where the first triplet in the random sequence is a stop codon. Overall, the 5,701 unique sequence clones code for 5,234 different peptides.

The dereplication algorithms used to generate the database provide information about the frequency of the different sequences in the library. The cluster size distribution is shown in suppl. Figure 1. It has a right skewed distribution (mean: 20,274, median: 9,870 sequences per cluster) with one extreme outlier with 4.2×10^7^ sequences, which corresponds to the vector plasmid without insert (“empty” vector).

The ORF length distribution in the database has the expected composition and features of a random database of sequences, i.e., it follows largely the expected distribution of predicted peptide lengths (suppl. Figure 2). Deviations concern mostly the longest sequence classes, due to the constant sequences derived from the vector. Note that some of the longest classes are also partly derived from frameshift versions.

With respect to GC content, we found that the sequences in the databases do not fully reflect a completely random synthesis. The mean and median GC content of the full reads is 53.8 %, and median GC content of the predicted ORFs is slightly higher (mean 53.04 %, median 54.6 %) with larger variance due to the shorter sequences (suppl. Figure 3A). A closer look to the GC content at every position in the database for reads with exactly the designed sequence length revealed a generalised bias towards lower A and higher G content at every position, remarkably larger on the 3’ end of the sequences starting at position 36 (suppl. Figure 3B). This is probably due to a bias during library synthesis, with a presumptive new supply of chemicals in between. Still, given that the length distribution of resulting peptides conforms mostly to the random expectation (compare suppl. Figure 2), we consider the library as being primarily made up of random nucleotide sequences.

A relevant descriptor of the structural properties of an amino-acid sequence is its intrinsic disorder level. Intrinsically disordered proteins lack defined secondary and tertiary structures, and naturally occurring genes have a higher intrinsic disorder than random sequences (Basile, Sachenkova, Light, & Elofsson, 2017; Heames, Schmitz, & Bornberg-Bauer, 2020; Wilson et al., 2017; Yu et al., 2016). In addition to intrinsic disorder scores, GC content (Basile et al., 2017) and amino acid content are used as indicators of the disorder levels of proteins in a database of sequences. Since large, hydrophobic amino acids are more likely to promote aggregation or formation of secondary structures, they are called order-inducing amino acids. The propensity of amino acids to induce order or disorder is one of the factors used for the calculation of intrinsic disorder scores (Campen et al., 2008).

Intrinsic disorder for the proteins in the database was calculated as the average intrinsic disorder score (IDS) of all residues in the peptide, using the -short setting of IUPred2A (see Methods). Average IDS values have a right-skewed bimodal distribution with the majority of sequences having an average IDS of 1.00 (suppl. Figure 4A). This is due to the large number of short sequences in the database that are very unlikely to be able to make any secondary structures and are also under the limit of detection of the software used. Grouping the sequences into length classes shows this effect clearly. The mean of the distribution of average IDS shifts to smaller values for longer peptide lengths, ranging from 0.947 for the shortest peptides with less than 10 residues, to 0.281 for those with 48 or more residues (suppl. Figure 4B). There is also a general correlation of IDS with length (suppl. Figure 4C), as well as with GC content (suppl. Figure 4D).

### Frequency changes of clones during the growth experiments

For all sequencing files from the experiments, 80-90 % of clean reads were successfully mapped to the database, allowing us to calculate frequency changes during the experiments. Raw count tables were used to do enrichment analyses using DESeq2. Although this algorithm was originally designed for the analysis of RNAseq sequencing data, it is also frequently used for the analysis of amplicon sequencing data. The assumption behind this is that the distribution of data in amplicon sequencing should follow a near-log normal distribution, with many low frequency counts and few high-frequency ones. The overall results of the DESeq2 analyses with respect to categorizing clones with positive (POS), negative (NEG) or non-significant (NS) changes are summarized in Table 1.

There is some variation between single experiments, especially with respect to the number of clones in the POS group. This is not directly related to the experiment type, i.e., the two experiments with the lowest fraction of POS clones (Exp1 and Exp7) have different cycle times (3h vs. 24h). Similarly, the range of log2-fold changes for individual clones varies considerably (Table 1). This suggests that even small variations in experimental conditions can lead to somewhat different outcomes. However, for all experiments there are always more NEG clones than POS clones. The average across all experiments shows 16 % POS clones, 36 % NEG clones and 48 % NS clones.

With the new pipeline, we could also trace the overall performance of the empty vector in the different experiments using the log2-fold change values. In experiments 1 to 3 it goes slightly up, in experiments 5 to 9 it goes slightly down, and in experiment 4 it has no significant change (Table 1). Only in experiment 5 the down trend was stronger than the average in this experiment. Note, however, that the DESeq2 normalization procedure penalizes against large count numbers in a way that could make negative trends stronger. Overall, we conclude from these data that the peptide and RNA expressed from the vector itself has no strong influence on growth.

We assessed also whether translation of the flanking sequences has a specific effect. The first 4 amino acids (MKLS) of the peptides are coded by the vector (see Figure 1). Of the clones that express only these first 4 amino acids due to a direct stop codon in the random sequence only 4 % are POS while 78 % are NEG, indicating a negative effect of this peptide compared to the overall clone performance (Table 1). Interestingly, this overall negative effect is relieved when only one or two additional amino acids are translated, with the percentage of NEG clones falling to 53 % and 38 % respectively (Table 1). From this analysis we conclude that the vector-derived, constant N-terminal amino acids of the peptides have an overall negative effect on growth, which can be overcome by additionally coded amino acids or the RNA sequence components in the clones.

The C-terminus of the full-length peptides is formed by 3 constant amino acids plus the 8 aminoacid FLAG tag sequence (see Figure 1). Of the different clones with this translated FLAG tag sequence, only 3 % are POS while 75 % are NEG across most experiments (Table 1), which would indicate a negative effect of this sequence. However, we find also the two frameshift translation versions of this sequence among the clones and both show a similar excess of NEG versus POS effects (Table 1). This suggests that it is not the FLAG tag sequence that acts negatively, but that longer peptides have generally a higher likelihood to be in the NEG group. This is also supported by the fact that peptides with a length of 48+ but without including any of the FLAG tag versions show a similar bias towards NEG (Table 1) (see also the further analysis of the length effects below).

### Length, GC content and amino acid composition dependence

For the further analysis, we have assigned each sequence in the database into the categories POS, NEG or NS, based on having consistent category assignments in the majority of experiments (see Methods). Since most sequences (> 95 %) fall consistently within one of these three categories (with the remainder being inconsistent and therefore not further analysed), one can compare whether peptides of different length are equally represented in each of these groups. Figure 2 shows that this is not the case. The fraction of NEG clones is particularly high for the shortest and the longest peptides. This is most likely caused by the negative effects of the vector derived parts of the sequence, as discussed above. The relative fraction of POS and NS clones is particularly high in the length classes between 8-20 amino acids.

The GC content distribution of ORFs in each of the groups is depicted in Figure 3. The peaks are similar for all three classes at about 57 % GC, slightly higher than the average for the whole library which is at 53 % GC. The POS and NS peptides show broader distributions than the NEG peptides, with a stronger shoulder towards lower GC contents. The NEG peptides show a second peak at 42 % GC, mostly driven by the negative effect of the shortest clones with only the vector-derived peptide (see above).

**Figure 3.**
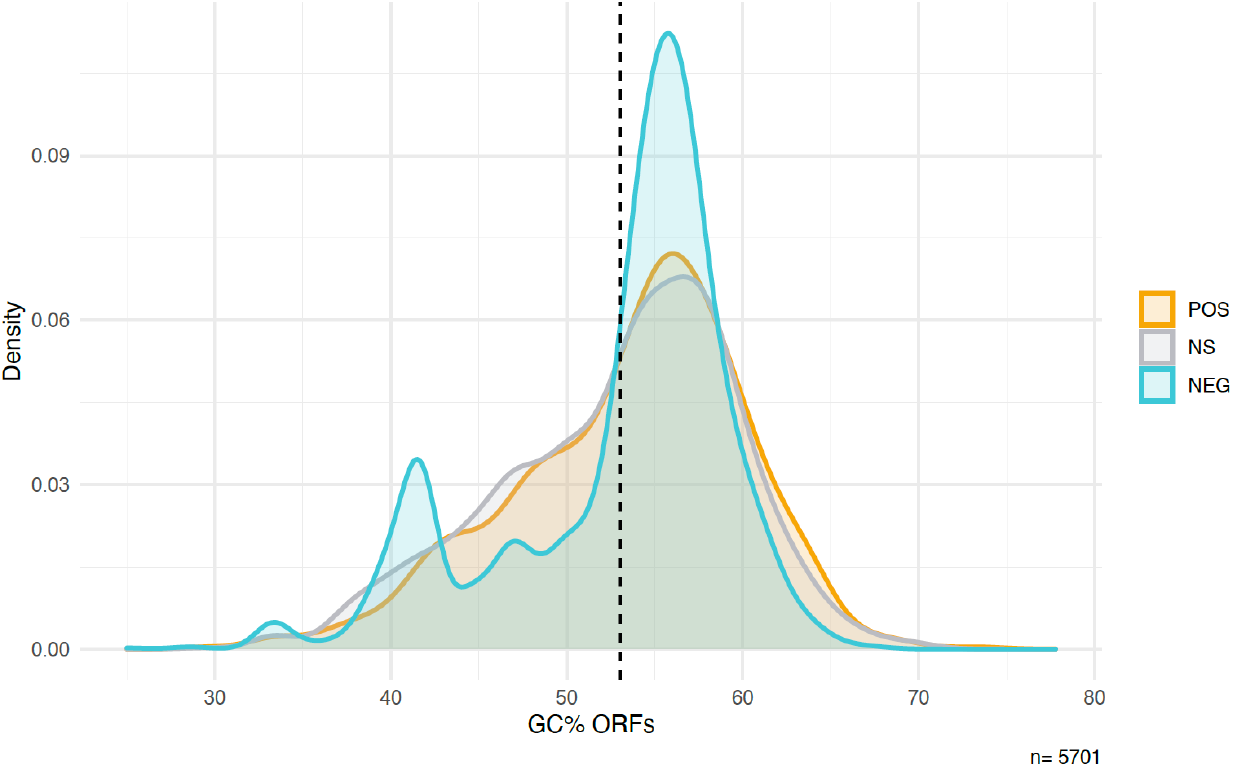
Density plots for the GC content of sequences in each of the three clone groups. Dashed line: average GC content of all ORFs in the library (53 %). The colours represent the assignment to the three groups of clones in the experiments: Light blue: sequences showing a decrease in frequency (NEG); dark grey: sequences showing no significant change in frequency (NS); orange: sequences showing an increase in frequency (POS).

We have also compared amino acid compositions of the peptides from the three clone groups. For this analysis we excluded the vector derived parts of the sequences. The overall frequencies for the whole database and the three groups of peptides are presented in suppl. Table 1. Figure 4 shows the differences for each group compared to the database. The largest differences are found for A, G and S in the comparison between POS and NEG groups. It is also notable that the frequency of 7 out of the 10 amino acids considered to be more disorder-inducing is lower in the NEG group than in the database, while 9 out of 10 of the order-inducing amino acids are depleted in the NS group. The POS group shows in general the largest deviations from the database.

**Figure 4.**
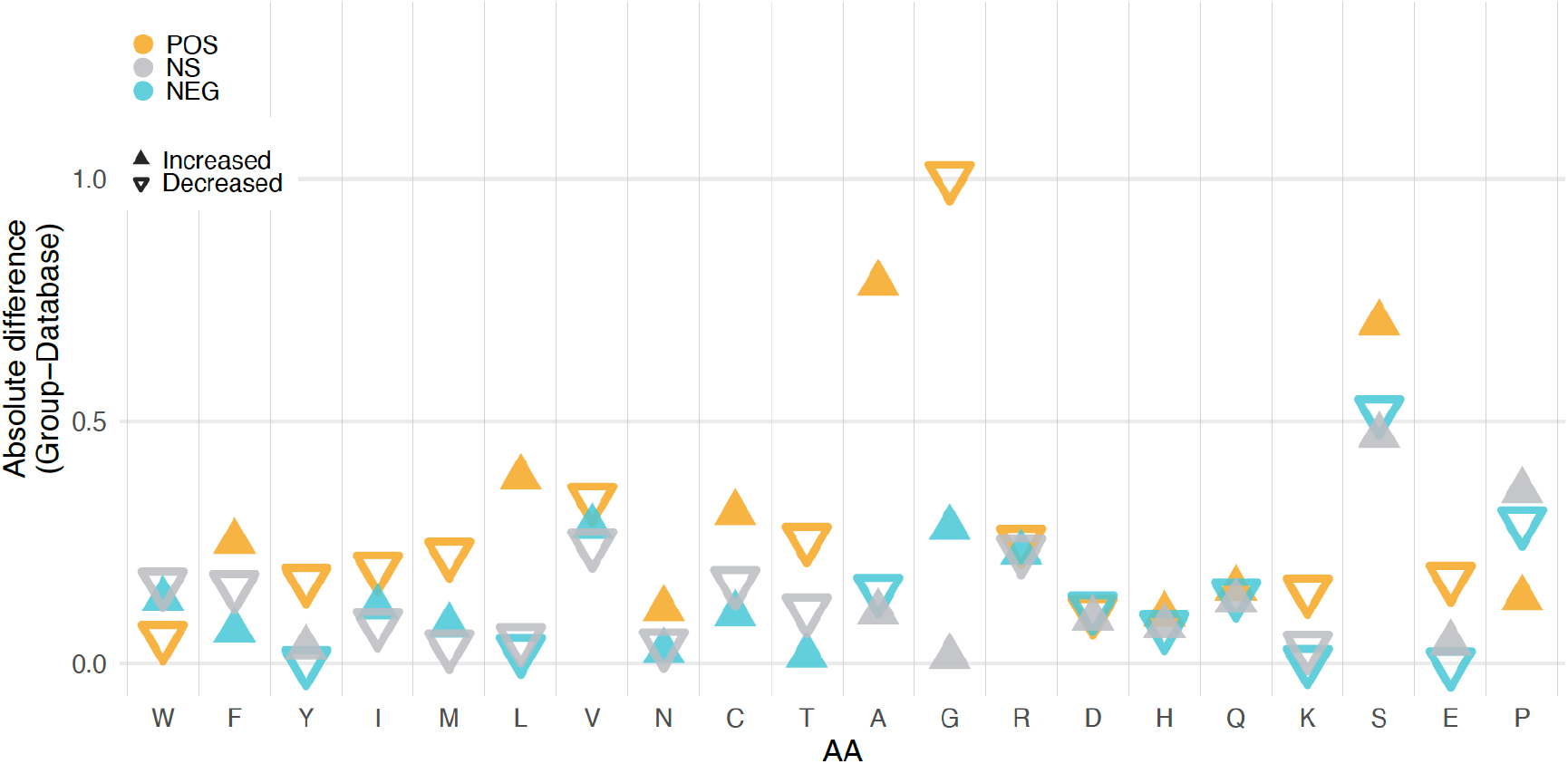
Differences in amino acid frequencies for the three groups of peptides. Frequencies were calculated as the percentage of each amino acid in all sequences in the groups and the complete database. Differences are shown as absolute values and the direction of the change is represented as +, if positive, or – is negative for each comparison. Light blue: NEG minus the database; grey: NS minus the database; orange: POS minus the database. Amino acids are ordered from left to right according to the TOP-IDP scale that reflects propensity for disorder induction from order promoting (left) to disorder promoting (right) (Campen et al., 2008).

### Structural features

The intrinsic disorder score (IDS) differs between the different clone groups, with the NEG group showing a stronger bimodal distribution than the two other groups (Figure 5). When breaking up the IDS in peptide length classes, it becomes clear that the highest IDS are due to the shortest classes (1-17 amino acids), for which the IDS calculation is anyway not very meaningful (compare also suppl. Figure 4). The lowest IDS scores are seen for the longest peptides (48+ amino acids), but otherwise there is no clear difference, especially between the POS and NS group of peptides (Figure 5).

**Figure 5.**
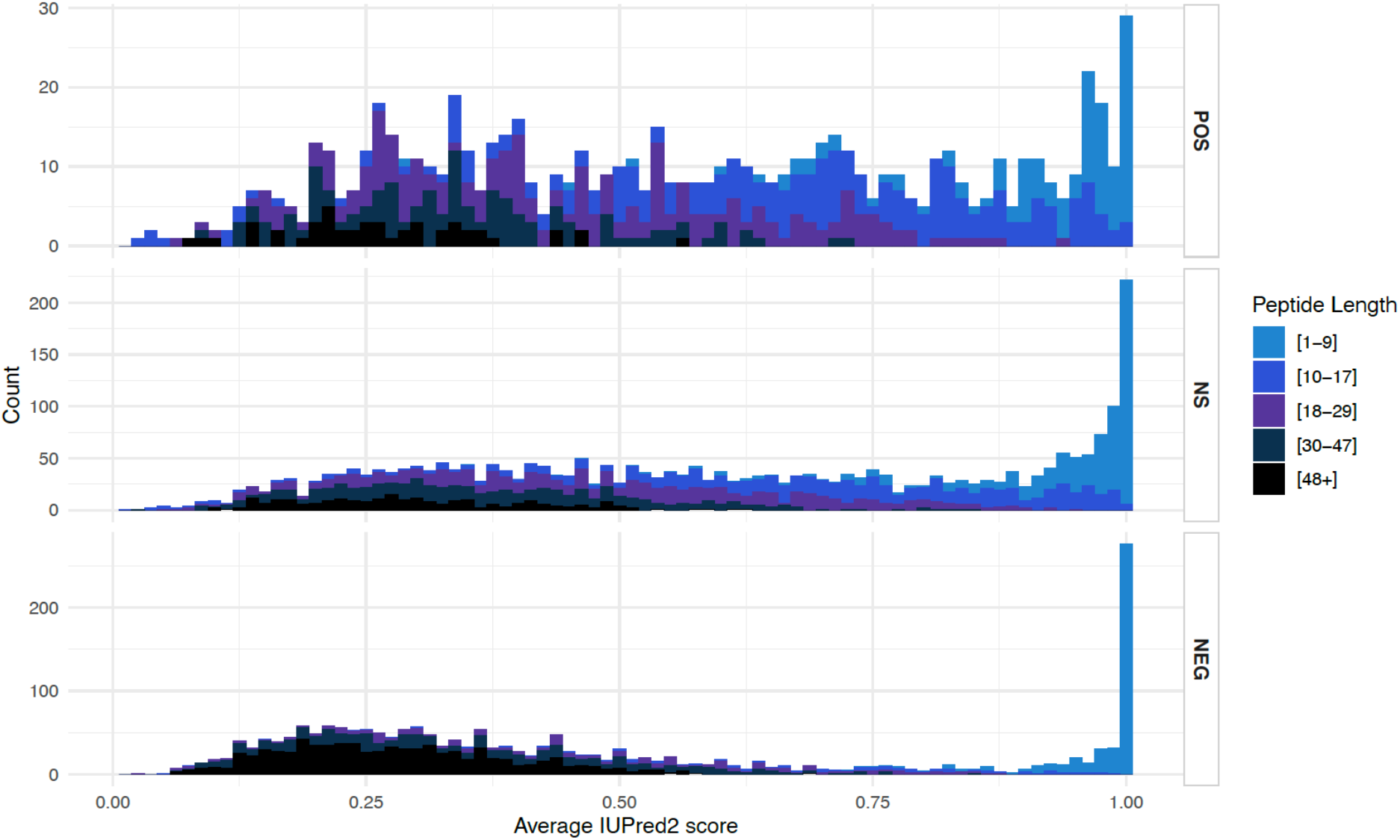
Intrinsic disorder scores (IDS) for the three groups of peptides. Histograms of average IUPred2 scores (IDS) coloured by the length categories depicted to the right. OR Empirical cumulative distribution of average IUPred2 scores (IDS) coloured by the length categories.

There are generally few highly ordered sequences in the library (i.e., sequences with an average IUPred2 score of less than 0.25). This could be due to the fact that highly ordered sequences tend to aggregate, and are expected to be highly insoluble and detrimental to the cells. In order to assess aggregation propensity, we used the software PASTA 2.0. It calculates the free energy of predicted ß-strand intermolecular pairings for each sequence and reports the lowest value for each peptide as the best pairing (I. Walsh et al., 2014). Lower aggregation energies mean that it is easier for the peptides to form amyloids or to aggregate. In general, aggregation energies lower than −5 pasta energy units (PEU) are considered evidence for possible amyloid formation. Sequences in the NEG group show generally lower PEU values than the two other groups with a peak at - 4 PEU and a distribution shifted towards even lower values (Figure 6A). There is also a secondary peak at aggregation energies higher than the other two groups (Figure 6A).

**Figure 6.**
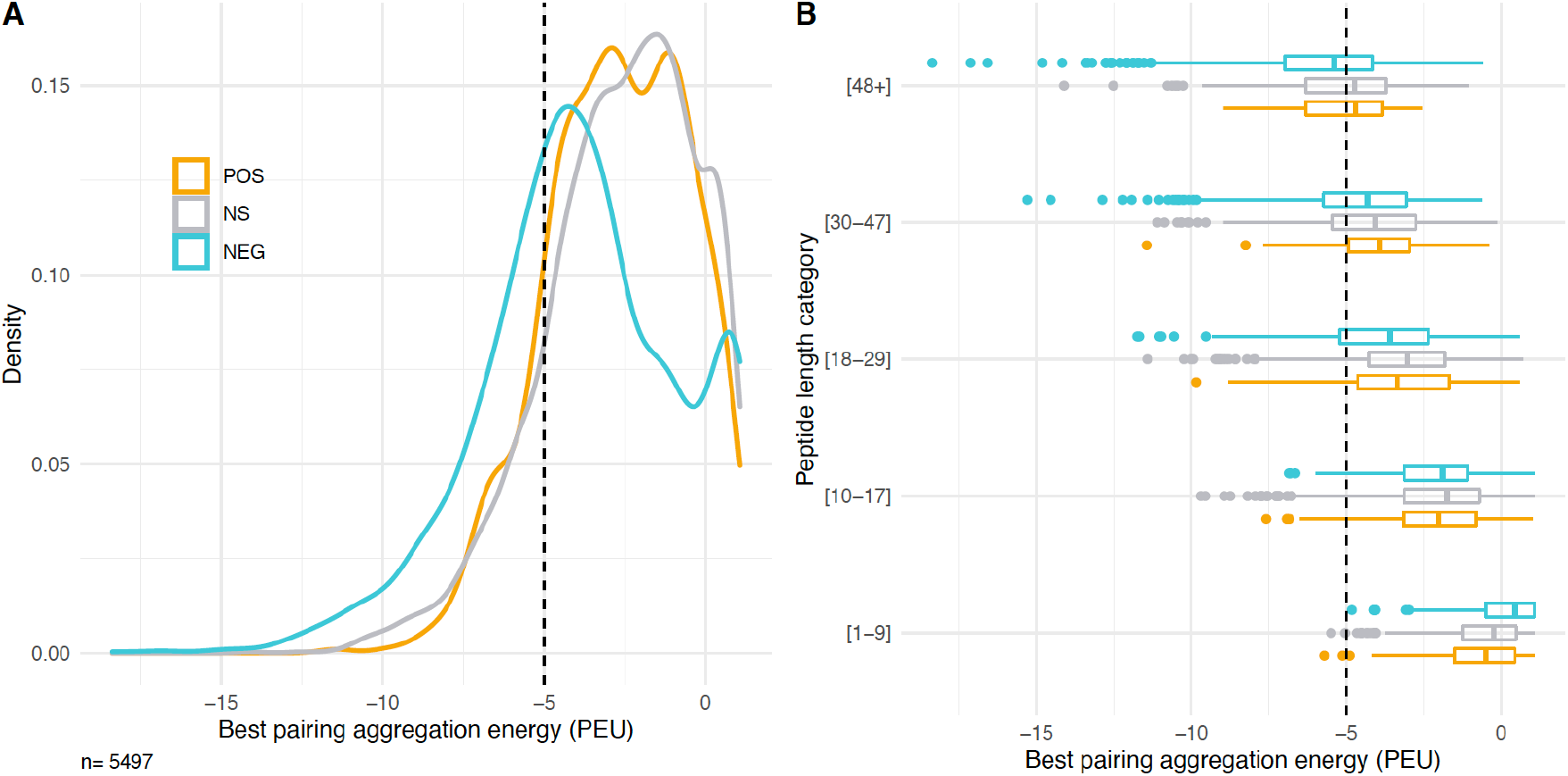
Aggregation energy analysis for the three clone groups and different length classes of peptides. (A) Density plots for the best aggregation energy of sequences in each group. (B) Best aggregation energy of sequences in each group and each length class. The lower and upper hinges correspond to the first and third quartiles (the 25th and 75th percentiles). The whiskers are the minimum and maximum data points up to 1.5 times the closest IQR.

In order to see whether sequences of a particular length are driving the observed pattern, we compared the data distribution for the different peptide length classes (Figure 6B). Interestingly, the distributions show different patterns between the groups of clones at the highest and lowest length values, but more similar ones at the intermediate ones. The secondary peak found in the NEG group at very high aggregation energies seems to be generated mostly by the shortest peptides, and the shift towards negative values, by the longest ones.

## 4. Discussion

Here we have performed an in-depth analysis of all available data from amplicon sequencing experiments of a library of *E. coli* cells expressing different randomly synthesized sequences and grown for four expansion cycles to allow competition between clones. The experiments, first described by (Neme et al., 2017), were set up to assess which fraction of random DNA sequences expressed in a living organism have the potential of producing molecules that have an effect on cell growth or fitness. This question is relevant for the study of the origin of innovation in biological systems and, in particular, of *de novo* genes derived from more or less random non-coding sequences.

The main goal of the present study was to broaden the analysis to all peptides in the data, irrespective of their length. (Neme et al., 2017) had originally focused on the full-length peptides only, with FLAG-tags derived from the vector sequences. The broadening of the focus to all expressed peptides allowed us to investigate whether the constant sequences flanking the random inserts in the library influenced the growth effects. In addition to this, we wanted to test whether there are particular molecular or structural features of the sequences driving their effect on the growth of the cells in this population. Studies looking at young and novel genes in diverse species report that novel genes and protogenes can have distinct features such as ORF length or intrinsic disorder levels that differentiate them from older genes and intergenic regions (Carvunis et al., 2012; Heames et al., 2020; James et al., 2021; Schmitz, Ullrich, & Bornberg-Bauer, 2018; Wilson et al., 2017; Yu et al., 2016). A possible explanation for these observations is that certain features make sequences more likely to be positively selected—or at least not selected against.

### New analysis pipeline

The first step in our analysis was to ask whether amino acid sequences of different lengths present in the population and not analysed in the original publication show a similar behaviour as the fulllength (65 amino-acid-long) sequences that were the focus of the (Neme et al., 2017) study. This required us to generate a new analysis pipeline which addresses three limitations of the original one. The first two—the incomplete removal of PCR and sequencing errors from the database, and the resulting artificial redundancy—are the result of using the predicted translation of the ORFs to generate the reference database of clones, instead of the nucleotide sequences. This was done to take advantage of the genetic code redundancy and to, at least partially, compensate for PCR and sequencing errors. The strategy, however, was insufficient as can be seen from the finding in the original publication of several very similar clones coding for peptides with only one or a few substitutions—an extremely improbable event in a library composed of random sequences of that length.

To compensate for such PCR and sequencing errors in this new analysis, we used a dereplication approach that removes all singleton reads from each sequencing file before joining them together and clustering them to a 97 % identity. We used the full-length merged and trimmed reads for database generation and mapping, which allowed us to keep track of all clones that code for the same (shorter) peptides independently, which was important to detect protein vs. RNA effects (see further discussion below). The database generated was composed of more than 5,000 reliably identifiable sequences predicted to code for peptides of all expected lengths. Furthermore, from its composition, it is possible to confirm that the library was indeed generated from random sequences, albeit with a slight bias towards a higher guanidine content during the synthesis process.

The final improvement to the pipeline was to change the algorithm used for mapping the reads back to the database from a local to a global alignment strategy. This greatly improved the speed and accuracy of the pipeline. Over 90 % of all reads containing the flanking sequences mapped back to our database of unique sequences in all experiment replicates. This represents a 30-50 % increase of mapped reads when compared with the pipeline used for analyses in the (Neme et al., 2017) study. As a result of this, we found that the change of frequencies can actually be much larger than what was initially reported. Some sequences, for example have a decrease in frequency between the first and the last cycle of the experiments of up to 1000-fold.

### Clone effects

Having identified how the frequency of sequences changes in the available experiments, we were able to classify the sequences in groups according to the direction of the change. With the improved mapping pipeline, we found that over 80 % of the sequences in the database had a consistent behaviour in at least 5 of the 9 experiments, suggesting that the observed results are indeed an effect of the sequences and not due to chance or drift. This is noteworthy, considering that the experiments were performed independently, by different researchers, at different times, and have variations between sampling schedules, seed size, sequencing depths and number of replicates.

Over half of the sequences in the database were consistently assigned to be either neutral (48 %) or to go up in frequency (16 %), suggesting that they are at least not very deleterious to the cells expressing them. This large proportion of sequences tolerated in a population of *E. coli* suggests that random sequences could be expressed and maintained in large numbers also in natural populations, making them an abundant source for possible evolutionary innovations.

We have also specifically evaluated the vector effects that were suggested to cause indirectly the observed positive effect (Knopp & Andersson, 2018; Weisman & Eddy, 2017). In dedicated experiments, (Knopp & Andersson, 2018) found that just expressing the 38 amino acid peptide from the empty vector (i.e. without a cloned insert), has a slightly negative effect on the exponential growth of the *E. coli* cells. By disrupting this vector peptide with a potentially neutral peptide, one could generate an apparent positive effect. However, we find in our analysis that this peptide behaves mostly like a neutral peptide in the context of the full experiment, i.e. when not only focussing on the exponential growth phase as done in (Knopp & Andersson, 2018), but taking all competition cycles into account. While there is, on average, a small negative effect across experiments, it is not strong enough to explain the growth of most POS clones as merely its relief. Hence, we conclude that the, in principle, justified reservations about positive effects in our experiments (Knopp & Andersson, 2018; Weisman & Eddy, 2017) are not warranted in the face of the full data shown here, as well as the arguments provided previously (Tautz & Neme, 2018).

### Negative effects of vector coded amino acids

Our data show that the first four amino acids expressed by the vector have by themselves a negative fitness effect on the cells. 73 % of clones encoding only the first 4 residues consistently decrease in frequency in 5 or more experiments. A reason for this might be that the second and third codons in the sequence—lysine (AAG) and leucine (CTT), respectively—are not the most commonly used by *E. coli* for these amino acids. Interestingly, this negative effect diminishes quickly when one or two additional amino acids are translated. Hence, it is not of much concern for the overall experiment with mostly longer peptides, although it contributes to the observed bimodal distributions of peptide length, intrinsic disorder and aggregation propensity for the NEG peptides.

The same observation demonstrates, however, that not only the coding part of a random sequence is important for its maintenance in a population. Over 20 % of the clones coding for the same peptide, but with different RNA sequences, show different growth trends in at least 5 experiments. In other words, clones with the same coding peptide had different effects on the growth trajectory of cells, due to the non-coding parts of their sequence. (Neme et al., 2017) had already shown that the RNA can have a different effect on growth than the protein by introducing a stop codon in single clones, disrupting the reading frame but keeping the rest of the sequence intact.

Systematic studies on replacing non-coding positions in an artificially expressed GFP RNA in *E. coli* have also shown that even small differences in RNA sequence can have differential fitness consequences for the cells (Mittal, Brindle, Stephen, Plotkin, & Kudla, 2018), although this might be mostly caused by perturbing co-translational protein folding (I. M. Walsh, Bowman, Santarriaga, Rodriguez, & Clark, 2020). On the other hand, transcription has also been shown to contribute strongly to the metabolic burden that is caused by overexpressing genes in *E. coli* (Z. P. Li & Rinas, 2020). It is thus expected that the clone effects that we find are a combination of effects from the expressed RNA and protein together.

### Protein structure correlations

Notwithstanding the possible fitness contribution of the RNA of the clones, we have analysed protein structural properties in the three different groups of peptides. The most compelling difference between clones with POS or NEG responses is their length. Shorter peptides in the length range of 8-20 aa are prevalent in the POS and NS groups, while longer ones are prevalent in the NEG group. While it is generally known that newly evolved genes are shorter than older genes (Carvunis et al., 2012; Neme & Tautz, 2013; Schmitz et al., 2018), the differences we observe here are at a much smaller scale than what is usually studied, since the ORF lengths of 4-65aa in our database are often not even annotated. Interestingly, in a study on the phenotypic impact of random sequences in Arabidopsis (Bao et al., 2017) used also very short peptides (with cores of six or 12 random amino acids) and found a substantial fraction having an effect on the phenotype, including possibly beneficial ones.

The NEG group of peptides shows in average lower intrinsic disorder and higher aggregation propensity compared to the POS group. This is in line with the observation that naturally occurring young genes are more likely to have higher intrinsic disorder (Wilson et al., 2017), which could be the reason why they are better tolerated by the cells (Tretyachenko et al., 2017).

We find no major differences in the three groups of peptides with respect to GC-content. But there are some differences with respect to overall amino acid composition. The largest contrasts occur between POS and NEG peptides, whereby POS peptides have more alanine and serine but less glycine. With its six codons, serine is a frequent amino acid in the random sequences and it has a strong disorder promoting effect (Campen et al., 2008). This could explain the higher disorder tendency in the POS peptides. Alanine and glycine, on the other hand, have both four codons and are therefore expected to occur equally frequently in random sequences and they have similar disorder promoting effects. It is therefore unclear why alanine is more prevalent in POS and glycine is more prevalent in NEG clones.

An additional possible implication of the enrichment of serine in both the POS and NS groups is its potential for evolution. Creixell et al. (Creixell, Schoof, Tan, & Linding, 2012) found that serine is the fastest-evolving amino acid and attribute this to fact that its six codons can be divided into two, very different, groups (AGY and TCN). The fact that the codons are so different facilitates non-synonymous substitutions, which allows evolution to explore a large sequence space in a shorter period of time. If, as our data seem to show, sequences containing larger fractions of serine are better tolerated by the cells, such sequences would be excellent starting material for the evolution of new functional peptides.

## 5. Conclusions

Although no single determining feature of a sequence could be identified that would earmark individual peptides as having potentially positive or negative effects on the cells, some differences exist with respect to structural properties. In particular, we found that shorter and more disordered peptides have a greater potential for being retained in a population as a primary source for novel genes, supporting the conclusions by James et al. on the general patterns of protein domain evolution (James et al., 2021). Most importantly, our data confirm that random sequences have the potential of being beneficial for the cell, especially in the context of the complex competition between clones that we study in these experiments. However, we show in the accompanying paper (Bhave & Tautz, 2021) that individual candidate POS clones can provide also a growth advantage in pairwise competition experiments, although not necessarily with the same strength as seen in the bulk experiments. We conclude that our experiments support the notion that random sequences are an abundant source for generating evolutionary novelty.

## Author Contributions

Conceptualization, Johana Fajardo and Diethard Tautz; Data curation, Johana Fajardo; Formal analysis, Johana Fajardo; Investigation, Johana Fajardo; Supervision, Diethard Tautz; Validation, Johana Fajardo and Diethard Tautz; Visualization, Johana Fajardo; Writing – original draft, Johana Fajardo; Writing – review & editing, Diethard Tautz.

## Data Availability Statement

Publicly available datasets were analyzed in this study. This data can be found here: Dryad http://dx.doi.org/10.5061/dryad.6f356 and at the European Nucleotide Archive (ENA) under the project number PRJEB19640. Sequencing data files for the replication experiments will be submitted to ENA, project number: ###. All codes for data organization and analysis are available as GitLab project under acc. number ###.

## Acknowledgments

We thank Ellen McConnell for her help in getting the replication experiment established and Rafik Neme for his comments and suggestions. The work was funded by institutional funds of the MPG to DT. JF is a member of the IMPRS for Evolutionary Biology.

