## Supplementary figures and tables for "The effects of sequence length and composition of random sequence peptides on the growth of *E. coli* cells"

### Supplementary files

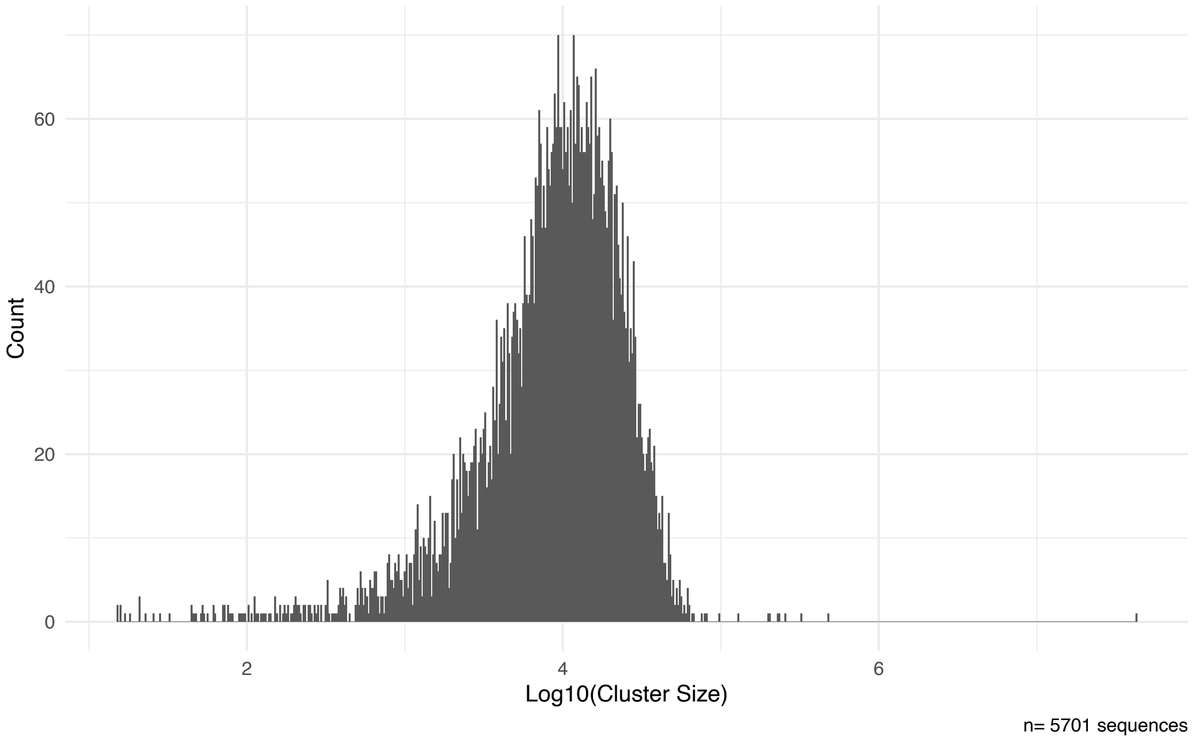


**Supplementary Figure 1**

Cluster size distribution of clones. Cluster sizes correspond to the number of reads across all experiments assigned to each of the 5,701 clones in the database. Most clones had between $10^3$ and $10^5$ reads assigned to them (mean $2.03 \times 10^4$ reads, median $9.9 \times 10^3$ reads). There is a single clone with $4.23 \times 10^7$ reads, which corresponds to the pFLAG-CTC plasmid without an insert.

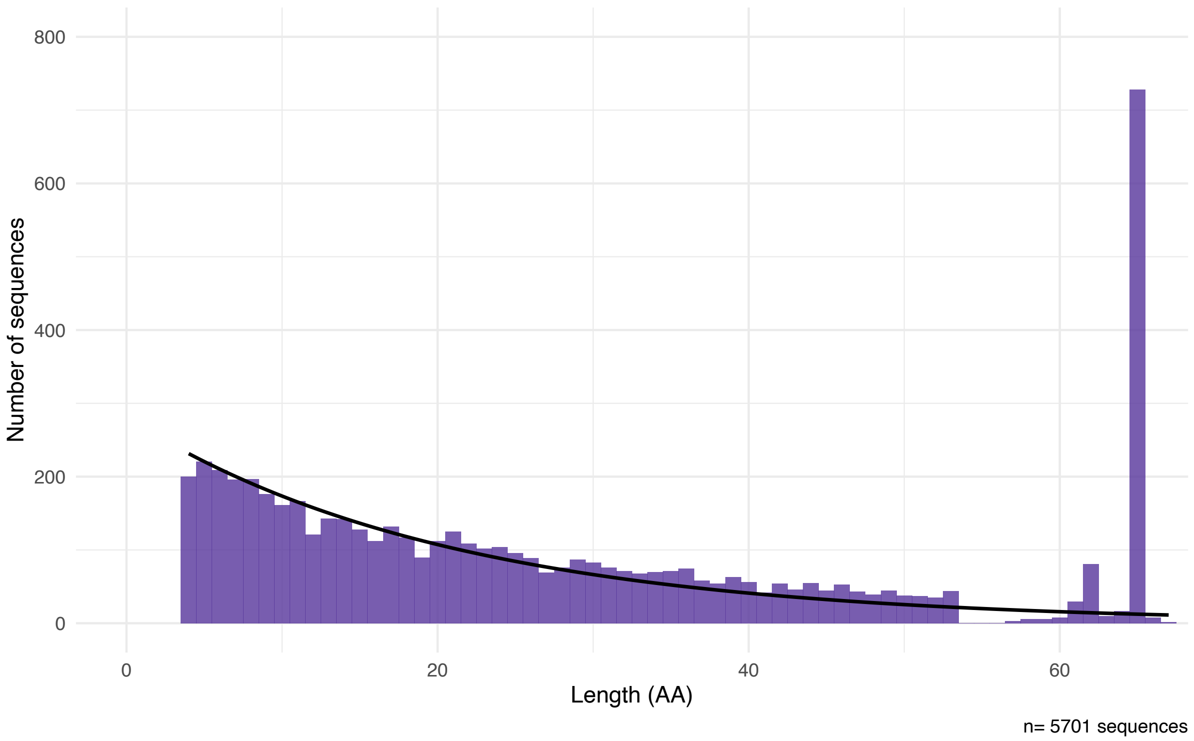


**Supplementary Figure 2**

Predicted peptide length distribution of all clones in the library. The library follows the expected distribution of stop codons in 5,701 random sequences, defined as the probability mass function of a geometric distribution with $p = 3/64$ stop codons (black line). Only the first ORF in each sequence was considered. Minimum peptide length corresponds to the 4 residues encoded by the plasmid and insert design (MKLS). Residues 55 to 65 are constant in the sequence design (ALVDYKDDDDK*), which should result in a single peak for sequences of length 65. However, the process of library synthesis or cloning generated 1735 clones with unexpected sequence lengths (30.43% of the clones in the library). This resulted in 171 clones with predicted peptide lengths between 55 and 64 residues or longer than 65 residues.

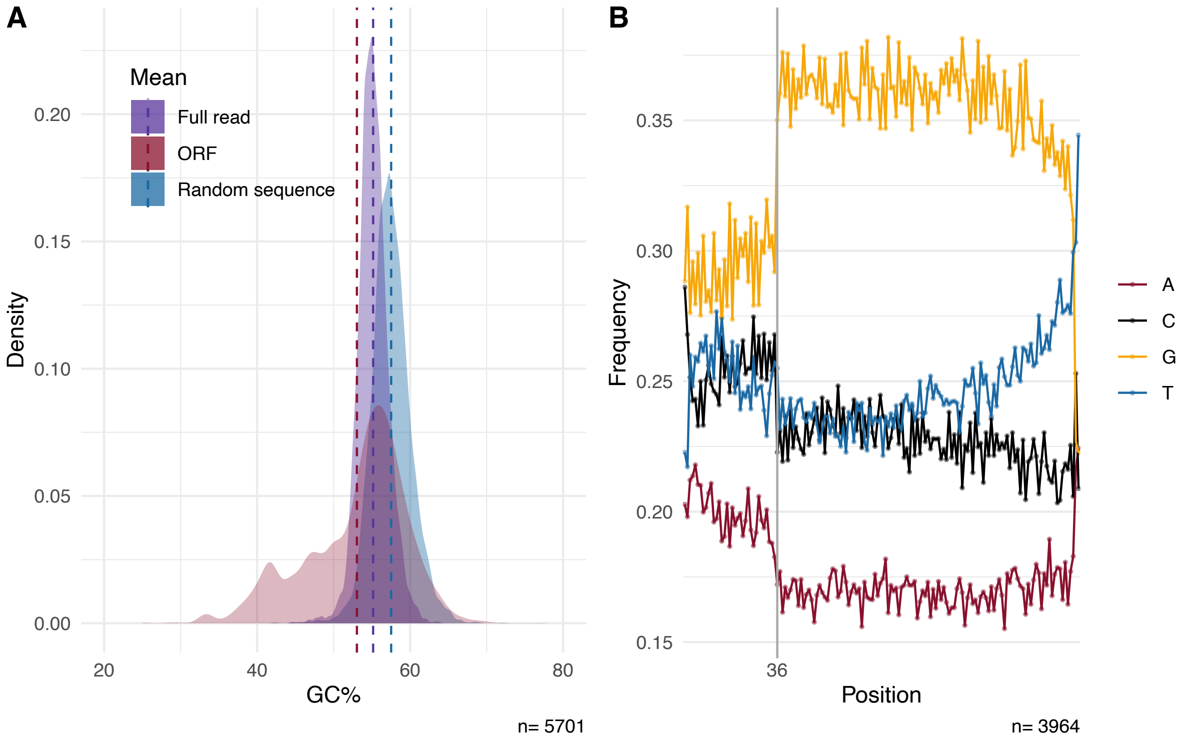


**Supplementary Figure 3:** GC content analysis of clones in the library.

A) GC content distributions calculated over the complete reads, between start and FLAG-tag sequences (this includes the flanking sequences of the plasmid, with a GC content of 46.2%the full sequenced reads (purple), the ORFs only (red) and simulated random sequences (blue). The ORF GC distribution is much more broadly spread, due to increased variance caused by very short ORFs.

B) Fraction of each nucleotide along the positions of the randomly synthesized sequence stretch.

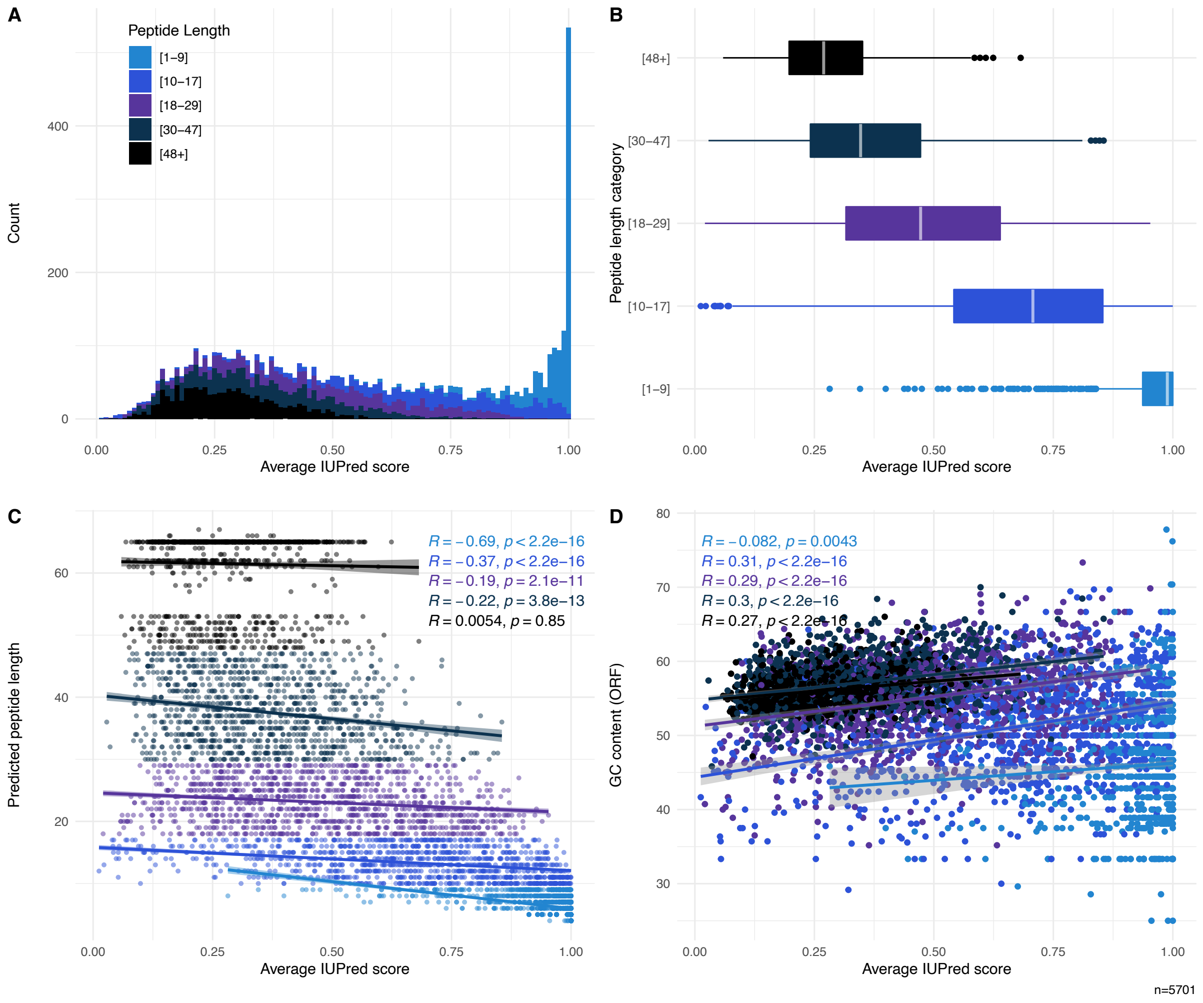


**Supplementary Figure 4**

Intrinsic disorder scores for peptides in the whole library in four different representations (A-D; see main text). Average intrinsic disorder scores become smaller for longer peptides (length 1-9 = 0.947, length 10-17 = 0.677, length 18-29 = 0.479, length 30-47 = 0.362, length 48+ = 0.281.

**Supplementary Table 1:** Amino acid frequencies for the database, as well as the three groups of peptides

| AA | Database | UP | DOWN | NS |
|---|---|---|---|---|
| W | 0.0284 | 0.0279 | 0.0298 | 0.0268 |
| F | 0.0320 | 0.0345 | 0.0326 | 0.0304 |
| Y | 0.0221 | 0.0204 | 0.0221 | 0.0224 |
| I | 0.0314 | 0.0294 | 0.0325 | 0.0306 |
| M | 0.0156 | 0.0134 | 0.0164 | 0.0153 |
| L | 0.0918 | 0.0957 | 0.0915 | 0.0913 |
| V | 0.0850 | 0.0816 | 0.0878 | 0.0825 |
| N | 0.0189 | 0.0200 | 0.0191 | 0.0185 |
| C | 0.0431 | 0.0462 | 0.0441 | 0.0414 |
| T | 0.0448 | 0.0422 | 0.0450 | 0.0437 |
| A | 0.0884 | 0.0963 | 0.0869 | 0.0895 |
| G | 0.1134 | 0.1034 | 0.1162 | 0.1136 |
| R | 0.1141 | 0.1116 | 0.1164 | 0.1118 |
| D | 0.0268 | 0.0257 | 0.0257 | 0.0277 |
| H | 0.0243 | 0.0253 | 0.0235 | 0.0251 |
| Q | 0.0264 | 0.0280 | 0.0250 | 0.0277 |
| K | 0.0205 | 0.0190 | 0.0204 | 0.0201 |
| S | 0.0904 | 0.0974 | 0.0852 | 0.0951 |
| E | 0.0282 | 0.0264 | 0.0282 | 0.0286 |
| P | 0.0545 | 0.0558 | 0.0516 | 0.0580 |

**Supplementary Table 2:** List of primers used

| | |
|---|---|
| pFLAG-CTC FWD-1 | AATGATACGGCGACCACCGAGATCTACAC AACCGCAT ACACTCTTTCCCTACACGACGCTCTTCCGATCT CATCATAACGGTTCTGGCAAATATTC |
| pFLAG-CTC FWD-2 | AATGATACGGCGACCACCGAGATCTACAC AAGGCCTT ACACTCTTTCCCTACACGACGCTCTTCCGATCT T CATCATAACGGTTCTGGCAAATATTC |
| pFLAG-CTC FWD-3 | AATGATACGGCGACCACCGAGATCTACAC AGAGTGTG ACACTCTTTCCCTACACGACGCTCTTCCGATCT GT CATCATAACGGTTCTGGCAAATATTC |
| pFLAG-CTC FWD-4 | AATGATACGGCGACCACCGAGATCTACAC CACAAGTC ACACTCTTTCCCTACACGACGCTCTTCCGATCT CGA CATCATAACGGTTCTGGCAAATATTC |
| pFLAG-CTC FWD-5 | AATGATACGGCGACCACCGAGATCTACAC CGTTCCTA ACACTCTTTCCCTACACGACGCTCTTCCGATCT ATGA CATCATAACGGTTCTGGCAAATATTC |
| pFLAG-CTC FWD-6 | AATGATACGGCGACCACCGAGATCTACAC GCTTGGAT ACACTCTTTCCCTACACGACGCTCTTCCGATCT TGCGA CATCATAACGGTTCTGGCAAATATTC |
| pFLAG-CTC FWD-7 | AATGATACGGCGACCACCGAGATCTACAC GTCAACAC ACACTCTTTCCCTACACGACGCTCTTCCGATCT GAGTGG CATCATAACGGTTCTGGCAAATATTC |

| | |
|---|---|
| pFLAG-CTC RWD-A | CAAGCAGAAGACGGCATACGAGAT AACCGGAA<br>GTGACTGGAGTTCAGACGTGTGCTCTTCCGATCT A CTGTATCAGGCTGAAAATCTTCT |
| pFLAG-CTC RWD-B | CAAGCAGAAGACGGCATACGAGAT AGAGTGAC<br>GTGACTGGAGTTCAGACGTGTGCTCTTCCGATCT TC CTGTATCAGGCTGAAAATCTTCT |
| pFLAG-CTC RWD-C | CAAGCAGAAGACGGCATACGAGAT CAACTGGT<br>GTGACTGGAGTTCAGACGTGTGCTCTTCCGATCT CTA CTGTATCAGGCTGAAAATCTTCT |
| pFLAG-CTC RWD-D | CAAGCAGAAGACGGCATACGAGAT CGTTCGTT<br>GTGACTGGAGTTCAGACGTGTGCTCTTCCGATCT GATA CTGTATCAGGCTGAAAATCTTCT |
| pFLAG-CTC RWD-E | CAAGCAGAAGACGGCATACGAGAT CTGTTCAC<br>GTGACTGGAGTTCAGACGTGTGCTCTTCCGATCT ACTCA CTGTATCAGGCTGAAAATCTTCT |
| pFLAG-CTC RWD-F | CAAGCAGAAGACGGCATACGAGAT GCTTGCAA<br>GTGACTGGAGTTCAGACGTGTGCTCTTCCGATCT TTCTCT CTGTATCAGGCTGAAAATCTTCT |
| pFLAG-CTC RWD-G | CAAGCAGAAGACGGCATACGAGAT GTCAACTG<br>GTGACTGGAGTTCAGACGTGTGCTCTTCCGATCT CACTTCT CTGTATCAGGCTGAAAATCTTCT |
